# Human methylome variation across Infinium 450K data on the Gene Expression Omnibus

**DOI:** 10.1101/2020.11.17.387548

**Authors:** Sean K. Maden, Reid F. Thompson, Kasper D. Hansen, Abhinav Nellore

## Abstract

While DNA methylation (DNAm) is the most-studied epigenetic mark, few recent studies probe the breadth of publicly available DNAm array samples. We collectively analyzed 35,360 Illumina Infinium HumanMethylation450K DNAm array samples published on the Gene Expression Omnibus (GEO). We learned a controlled vocabulary of sample labels by applying regular expressions to metadata and used existing models to predict various sample properties including epigenetic age. We found approximately two-thirds of samples were from blood, one-quarter were from brain, and one-third were from cancer patients. 19% of samples failed at least one of Illumina’s 17 prescribed quality assessments; signal distributions across samples suggest modifying manufacturer-recommended thresholds for failure would make these assessments more informative. We further analyzed DNAm variances in seven tissues (adipose, nasal, blood, brain, buccal, sperm, and liver) and characterized specific probes distinguishing them. Finally, we compiled DNAm array data and metadata, including our learned and predicted sample labels, into database files accessible via the `recountmethylation` R/Bioconductor companion package. Its vignettes walk the user through some analyses contained in this paper.

## Introduction

DNA methylation (DNAm) has been widely studied for its roles in normal tissue development [1–4], biological aging [5–7], and disease [8–12]. DNAm regulates gene expression, either in *cis* if it occurs in a gene’s promoter, or in *trans* if it overlaps an enhancer or insulator [4,9,13]. Whole-genome DNAm (or “methylome”) analysis, especially in epigenome-wide association studies (EWAS), is a common strategy to identify epigenetic biomarkers with potential for clinical applications such as in prognostic or diagnostic panels [14–16].

Most investigations probe DNAm with array-based platforms. Published DNAm array data and sample metadata are commonly available through several public resources. These include cross-study databases like the Gene Expression Omnibus (GEO) [17,18] and ArrayExpress [19], as well as landmark consortium studies like the Cancer Genome Atlas (TCGA) [20] and the Encyclopedia of DNA Elements (ENCODE) [21,22]. Recently published databases and interfaces provide access to samples from these sources [23–27].

While over 1,604 DNAm array studies and over 104,000 samples have been submitted to GEO since 2009 (Figure S1), there have been few attempts to rigorously characterize technical and biological variation across studies. In 2013, two studies independently compiled DNAm array samples from GEO and elsewhere, analyzing epigenetic age across tissues and diseases [5], and investigating cross-study normalization [28]. More recently, a study from 2018 evaluated metadata and sample quality across 8,327 DNAm array samples [29].

Further, while the GEO website provides access to submitted experiment and sample metadata, the metadata are not necessarily structured and require harmonization to facilitate cross-study analyses. There are currently no R/Bioconductor [30] packages providing access to uniformly normalized array data across GEO studies accompanied by harmonized metadata. It should also be noted that most GEO studies do not include raw intensity data (IDAT) files, which are needed to uniformly normalize samples and thus limits their utility for novel cross-study analyses.

The vast majority of GEO DNAm array data is composed of samples using Illumina’s HumanMethylation 450K (HM450K) BeadArray platform. Restricting attention to HM450K samples with IDATs published on or before March 31, 2019, we identified 35,360 samples from 362 studies, over three times the number of samples studied by either [5] or [28]. From sample IDATs, we extracted raw signals and probe significance data, derived quality metrics from control probe data, and performed normalization on out-of-band signal with the noob method [31]. We also learned a controlled vocabulary of sample labels by applying regular expressions to metadata and used existing DNAm array-based models to predict sex, epigenetic age, and blood cell fractions [5,32,33]. We conducted analyses investigating the performances of standard quality assessments and identified studies with frequent failed samples. Finally, we characterized autosomal DNAm variation in 7,484 samples from seven non-cancer tissue types.

To aid other investigators interested in reanalyzing DNAm array data from GEO, we compiled raw and noob-normalized DNAm array data with our learned and predicted metadata into HDF5-based databases accessible using `recountmethylation`, a companion R/Bioconductor [30] package available at https://doi.org/doi:lO.l8l29/B9.bioc.recountmethylation. Use of this package is covered thoroughly in accompanying vignettes, which also reproduce some of the results contained in this paper.

## Results

### Recent growth in GEO DNAm array samples is linear

We obtained sample IDATs and metadata for studies from the Gene Expression Omnibus (GEO). GEO is the largest public database for human DNAm array studies, and the majority of GEO’s DNAm array samples use one of three of Illumina’s BeadArray platforms: the HumanMethylation27K (HM27K), the HumanMethylation450K (HM450K), and EPIC, also known as the HumanMethylation850K (HM850K). On GEO, we identified 104,746 unique sample accession numbers (GSMs) from 1,605 studies (GSEs) published using one of the three major Illumina DNAm array platforms (Figures 1a and S1). Among 1,605 published studies, 74% used HM450K, 21% used HM27K, and 5% used EPIC. Among 104,746 published samples, 79% were on HM450K, 18% on HM27K, and 3% on EPIC. All three platforms showed increasing publication rates of samples and studies over the first three years of their availability. Few new studies and samples from 2013-2018 used the HM27K platform, while samples and studies using HM450K have grown linearly through 2018.

**Figure 1.**
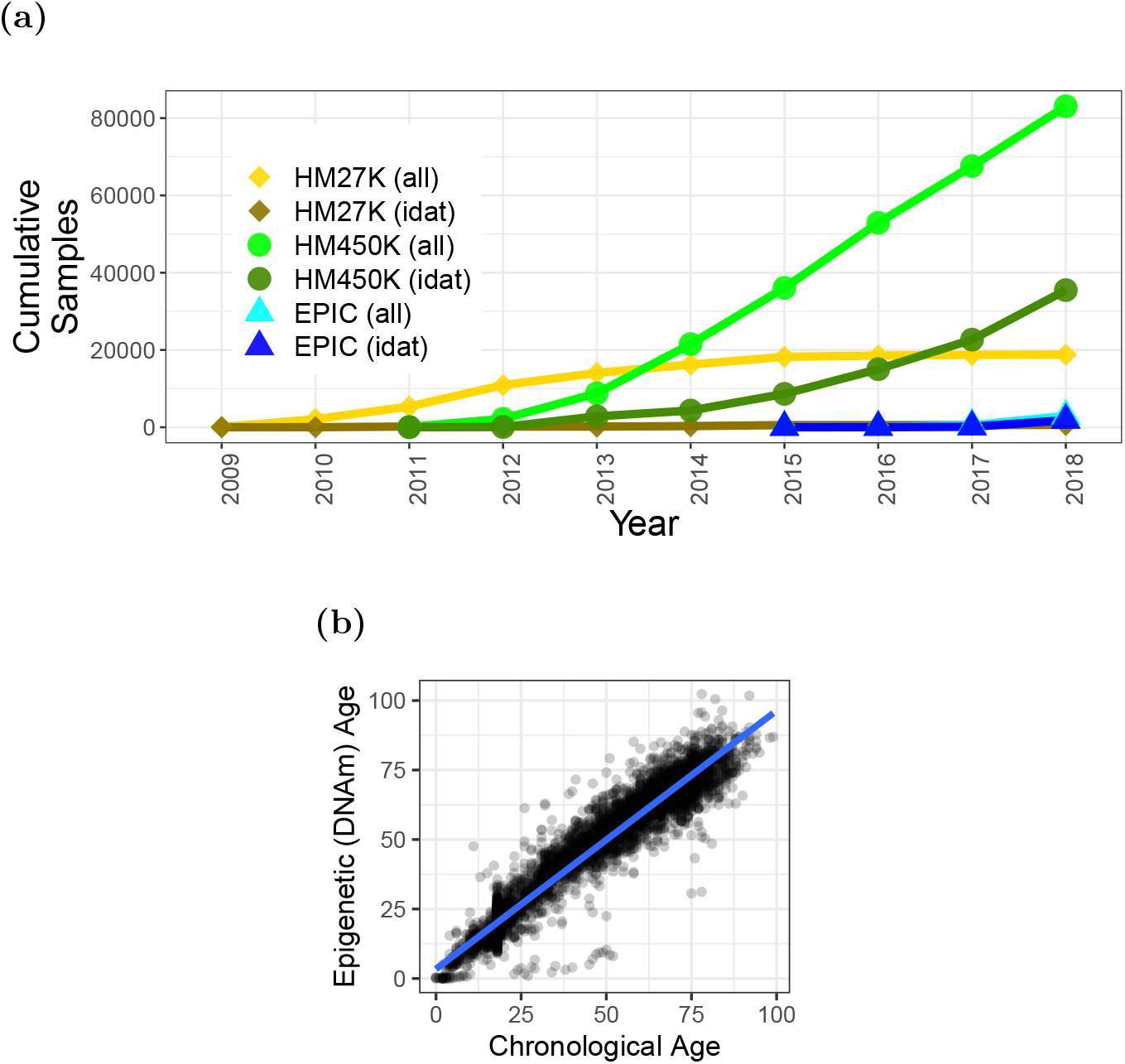
Cross-study summaries of DNAm array samples from GEO. (a) Cumulative samples by year using one of three major Illumina BeadArray DNAm array platforms (HM27K, HM450K, and EPIC/HM850K, point shapes), showing either all samples or subsets with available IDAT files for each platform (line colors). Samples with IDATs using the HM450K platform (dark green line, circle shape) were compiled and analyzed (see Methods and Results). (b) Scatter plot of chronological (x-axis) and epigenetic (y-axis) ages, with linear model fit (blue line), for 6,019 non-cancer tissues run using the HM450K platform. Chronological age was mined from GSE metadata. Epigenetic age was calculated using the model in [5] (Methods).

### Fewer than half of DNAm array studies on GEO include raw data

Raw data for a DNAm array sample is comprised of two IDAT files, one for each of the red and green color channels. Accessible raw data is important for uniform normalization of samples across studies, yet not all samples on GEO come with these data. In total, 37,919 samples (36% of total) included sample IDATs, where 93% were run on HM450K, 5% on EPIC, and 2% on HM27K. By platform, EPIC included the largest percentage of GSMs with available IDATs at 63%, followed by HM450K at 43%, and HM27K at just 3%. The more frequent availability of IDATs for newer arrays seems to reflect a significant shift in data submission norms well after the inception of the HM27K platform.

### Most annotated GEO HM450K samples with available raw data are from blood or brain

There were enough study and sample metadata for us to annotate 27,027 samples, 76% of the 35,360 we analyzed. We annotated these samples by applying regular expressions to the mined metadata. Our vocabulary for annotations was composed of 72 distinct terms (see Methods) strongly inspired by those used in the methylation array resource Marmal-aid [28]. Tissue terms for blood accounted for the majority (18,212 samples, 67% of total), followed by brain (6,690 samples, 25% of total), tumor (1,977 samples, 7% of total), breast (1,525 samples, 6% of total), and placenta (1,338 samples, 5% of total). We further annotated disease and experiment group for 22,790 samples (64% of total) using 38 distinct disease-and group-related terms. Among these, disease terms for cancer were assigned to over half (13,131, 58% of total) of samples, while terms for normal, control, or healthy were assigned to 10,808 samples (47% of total). The most frequently annotated cancers included leukemia (2,585 samples, 20% of total), breast cancer (511 samples, 4% of total), colorectal cancer (314 samples, 2% of total), and prostate cancer (196 samples, 1% of total). We compared disease and tissue characteristics to distinguish between tumor and normal samples from cancer patients, estimating that a third of samples were from tumor (see Methods and Table S1).

### Chronological age is accurately predicted from epigenetic age in non-cancer tissues

Prior work has shown chronological age can be predicted with high accuracy from DNAm among non-cancer tissues [5,6]. We used the epigenetic clock published in [5] to calculate an epigenetic age for each of 35,360 samples (Table S1). In 16,510 samples, analysis of variance (ANOVA) showed most epigenetic age variation was explained by chronological age (52% of variances, p < 2.2e-16), followed by experimental batch (i.e. GSE; 24%, p < 2.2e-16), cancer status (7e-2%, p = 1.3e-9), and predicted sample type (8e-3%, p = 1.6e-2). Between mined and predicted ages in these samples, high differences (12.9 years mean absolute difference, or MAD) and errors (R-squared = 0.76) likely resulted from study-specific conditions such as metadata inaccuracies or age label misattributions from our mining strategy as well as our inclusion of cancers and non-tissue samples (see Methods). In 6,019 likely non-cancer tissue samples across 37 GSE records with GSE-wise MADs ≤ 10 years, epigenetic age variance contribution from chronological age increased to 93% (ANOVA, p < 2.2e-16) and contribution from GSE decreased to 2% (p < 2.2e-16, Figure 1b). Unsurprisingly, the non-cancer tissue samples showed lower age differences (MAD = 4.5 years) and errors (R-squared = 0.94), and ages were highly correlated (Spearman Rho = 0.96, p < 2.2e-16), supporting the well-established finding that chronological age is accurately predicted from epigenetic age in non-cancer tissues [5, 6]. We imputed missing ages using epigenetic age for non-cancer tissue analyses (see Methods and Results below).

We next studied age acceleration [5,6] by probing the difference between epigenetic age and chronological age. Among the 68 samples we identified with outlying (≥ 15 years) positive age acceleration, the most frequently represented study was GSE61454 [34], which accounted for 18 adipose samples from severely obese patients. We observed 86 negative age acceleration outliers (≤ −15 years), including 14 saliva samples from control subjects in a study of Parkinson’s disease, GSE111223 [35], and 19 whole blood samples from CHARGE or Kabuki syndrome patients from GSE97362 [36]. In the latter study, we suspect reported ages are inaccurate and older than actual ages (private correspondence, investigation ongoing).

### Almost a fifth of samples fail at least one of 17 BeadArray quality control assessments

Illumina prescribes 17 quality assessments for its 450K array, each measuring the performance of a different step in a methylation assay such as extension or hybridization [37,38]. A given assessment comprises a quality metric and a minimum threshold value below which the assessment is failed. We call these assessments BeadArray controls. We used the 17 BeadArray controls and their minimum quality thresholds to evaluate assay qualities in 35,360 samples (Tables S2 and S3). Results are summarized in Figure 2a. The highest proportions of samples failed the non-polymorphic green and biotin staining green controls, with about 6.7% failing each (2,381 and 2,368 samples, respectively). By contrast, there are six BeadArray controls, each failed by fewer than 100 samples. A substantial number of samples (6,813, 19% of total) failed at least one control. Of samples that failed at least one control, 4,456 samples (66%) failed exactly one control, while 2,357 samples (34%) failed more than one control. Of samples that failed more than one control, 634 failed both biotin staining controls and 648 failed both non-polymorphic controls. Samples failing at least one control were significantly enriched for certain labels including “cord blood,” “brain cancer”, “prostate cancer”, “arthritis”, and “obese” (binomial test, BH-adjusted p < 1e-3).

**Figure 2.**
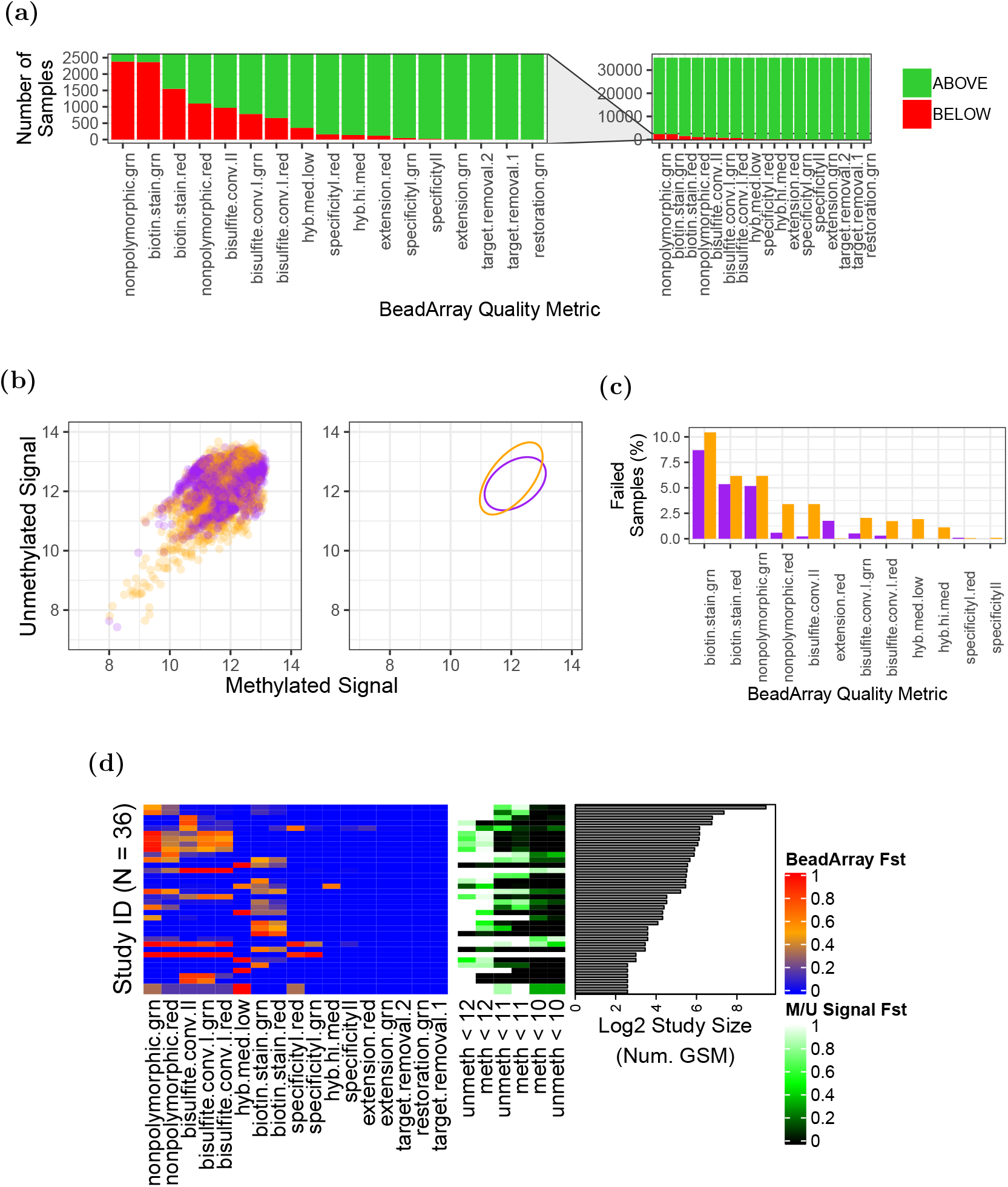
Quality analyses across samples, storage conditions, and studies. (a) Barplots counting samples (y-axis) falling above (light green) or below (red) manufacturer-prescribed thresholds across the 17 BeadArray controls (x-axis). Full view is on right, and magnification is on left. (b) Scatter plots (left) and 95% confidence intervals (right) for *log_2_* median methylated (x-axis) and *log_2_* median unmethylated (y-axis) signal of 3,467 FFPE (orange) and 5,729 fresh frozen samples (purple). (c) Percentages of FFPE (orange) and fresh frozen (purple) samples failing BeadArray controls. (d) Heatmaps depicting fraction *(Fst* in legends) of samples in a study failing quality assessments across 36 studies with high failure rates. BeadArray controls are shown on left, while methylated (M, meth) and unmethylated (U, unmeth) are shown in the middle. log_2_ study sizes are shown on the right.

### The intrinsic dimension of the 17 BeadArray controls is small

We studied signals and outcomes to determine how best to use the BeadArray controls for sample quality assessments. Cross-sample signal distributions for five BeadArray controls were bimodal, with distinct low- and high-signal modes; minimum quality thresholds fell near low-signal modes (Figure S3). For these controls, modifying minimum thresholds to more robustly capture low-signal samples could improve their utility. Principal component analysis (PCA) of sample control performances showed the top five components explained 84% of overall variances. Component-wise ANOVAs showed that just five of the 17 controls explained the majority of sum of squared variances across these top five components (minimum = 67%, maximum = 99%, median = 97%). This suggests the intrinsic dimensions of sample quality outcomes is much less than seventeen, and likely as small as five. These explanatory controls included both biotin staining controls, both non-polymorphic controls, and bisulfite conversion I red (Figure S4).

### FFPE samples fail at least one BeadArray control almost twice as often as fresh frozen samples

We investigated the impact of storage conditions on sample quality across 28 studies by comparing 3,467 formalin-fixed paraffin embedded (FFPE) and 5,729 fresh frozen (FF) samples (Table S4 and Figure 2b). FFPE samples showed greater variance than FF samples in both methylated (0.36 for FFPE vs. 0.27 for FF) and unmethylated (0.50 for FFPE vs. 0.21 for FF) signal channels. The trend could be driven either by condition-related sample characteristics (e.g. increased DNA deamination and/or lower DNA yield in FFPE, etc.) or differing preparation protocols (e.g., addition of the DNA restoration step for FFPE, [39–41]). Enriched labels also varied by storage condition among low-signal samples (binomial test, BH-adjusted p < 1e-3), where “colorectal,” “intestine,” and “mucosa” were enriched among FFPE, while “nasal,” “pancreas,” and “epithelial” were enriched among FF samples.

Across the 12 of 17 total BeadArray controls each with at least one failing sample, 228 FFPE samples (8.31% of total FFPE sample count) and 241 FF samples (4.21% of total FF sample count) failed at least one control. These failure rates are much lower than the failure rate across all samples (19%) and point to a correlation between study metadata completeness (e.g., inclusion of storage procedure details) and sample quality. FFPE samples failed 10 of the 12 metrics between 0.1-3.2% more often than FF, including all three bisulfite conversion metrics (Figure 2c). FF samples had higher failure rates for two BeadArray controls (extension red and specificity I red; see Table S4). While no samples failed the restoration BeadArray control, increasing the minimum threshold for failure from the default manufacturer-prescribed value of 0 to 1, which is recommended as an alternative in Illumina documentation [37,38], failed 69 FFPE samples (2%) and one FF sample (2e-3%). In summary, while FFPE samples were of lower quality than FF samples across assessments, the differences were modest, and the vast majority of FFPE samples passed all controls considered.

### 10% of studies each have >60% samples failing quality assessments

Across 362 studies, we evaluated the fraction *f_st_* of failed samples per study, defining a failed sample as one that either (i) fails at least one BeadArray control, or (ii) has log_2_ median methylated and log_2_ median unmethylated signals each <11 as described in [32] (see Table S5 and Figure S5). Of the 36 studies each with *f_st_* > 60% (Figure 2d), samples fail in each of 23 studies due only to (i), samples fail in each of five studies due only to (ii), and samples fail in each of the remaining eight studies due to either (i) or (ii). These 36 studies ranged in size from 6 to 692 samples and comprised a total of 2,020 samples, with a median study size of 23 samples. One study was of condition-specific DNAm data reliability (GSE59038, [40]) and included several stress tests of the assay, so many failed samples are not unexpected. Another study was GSE62219 [42] and included blood from 10 young individuals. We further noted the previous study [43] also determined these samples were of low quality.

### DNAm principal component analysis shows clustering by tissue with greater variances among cancers

We studied DNAm variance using PCAs of autosomal DNAm (Figure 3) as measured by noob-normalized Beta-values (see Methods). The first two components from PCA of 35,360 samples explained 35% of total variance, with PC1’s contribution 25% and PC2’s contribution 10% (see Figure 3a). Four outlying blood samples (PC1 > −10) included two from whole blood, one of T-cells, and one stem cell sample from umbilical cord blood (left plot of Figure S6a). For the top two components, leukemia samples showed greater variances than blood samples: the ratio of variance in PC1 for leukemia samples to variance in PC1 for blood samples was 1.25 (F-test p < 1e-2), and the ratio of variance in PC2 for leukemia samples to variance in PC2 in blood samples was 6.18 (F-test p < 1e-2). This is consistent with how (1) leukemia samples have greater variance than blood samples at each of the majority of probes (311,127, or 66%), and (2) leukemia samples have greater median variance than blood samples across probes (median Beta-value variance for blood samples = 1e-3, median Beta-value variance for leukemia samples = 5e-3; Figure S6b).

**Figure 3.**
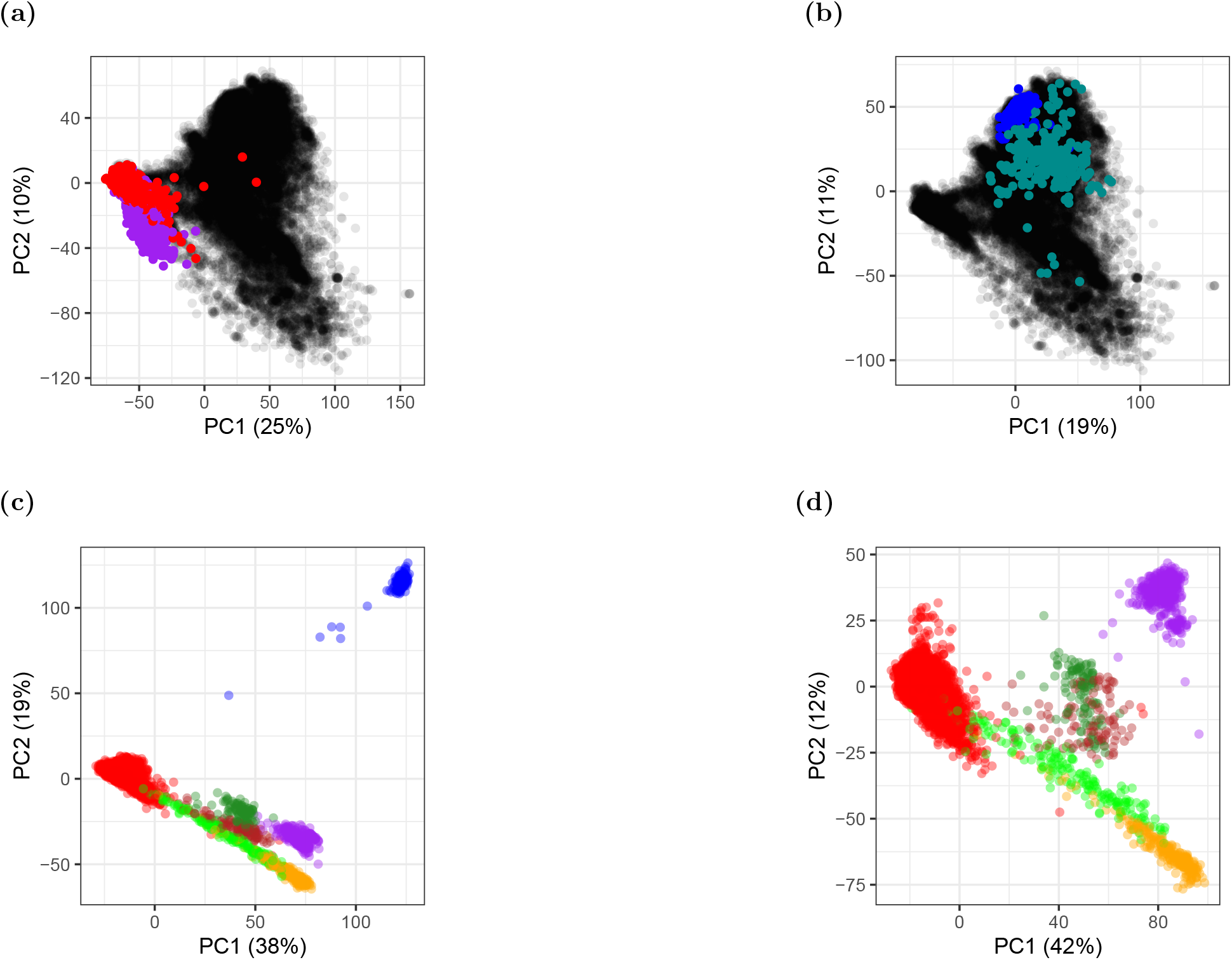
Scatter plots of top 2 components from principal component analyses (PCAs) of autosomal DNAm (Methods). Each axis label also contains percent of total variance explained by the component. (a) PCA of 35,360 samples, with color labels for non-cancer blood *(N* = 6,001 samples, red points) and leukemias (780, purple), and remaining samples (28,579, black). (b) PCA of 28,579 samples remaining after exclusion of blood and leukemias from (a), highlighting non-cancer brain (*N* = 602 samples, blue), brain tumors (221, dark cyan), and remaining samples (27,756, black points). Facet plots of sample subsets in (a) and (b) are shown in Figure S6. (c) PCA of 7,484 samples from 7 tissue types, including sperm (*N* = 230 samples, blue), adipose (104, dark red), blood (6,001, red), brain (602, purple), buccal (244, orange), nasal (191, light green), and liver (112, dark green). (d) PCA of 7,254 non-cancer tissue samples remaining from (c) after exclusion of sperm, with color labels as in (c).

From PCA of the 28,579 samples remaining after blood and leukemia samples were removed (Figure 3b), the first two components explained 30% of total variance, with PC1’s contribution 19% and PC2’s contribution 11%. Seven outlying (PC1 > 0, PC2 < −5) brain tumor samples included two primary tumors and one metastasis each from medulloblastoma cases, as well as four brain metastases from uncertain primary tumors, from the studies GSE108576 [44,45] and GSE63669 [46] (Figure S6c). For the top two components, brain tumors showed greater variances than non-cancer brain samples: the ratio of variance in PC1 for brain tumors to variance in PC1 for non-cancer brain samples was 12.05 (F-test p < 1e-5), and the ratio of variance in PC2 for brain tumors to the variance in PC2 for non-cancer brain samples was 22.40 (F-test p < 1e-5). This is consistent with how (1) brain tumors have greater variance than non-cancer brain samples at each of the majority of probes (444,304, or 94%), and (2) brain tumors have greater median variance than non-cancer brain samples across probes (median Beta-value variance for non-cancer brain samples = 1e-3, median Beta-value variance for brain tumors = 1e-2; Figure S6d). Our findings are consistent with previous evidence of higher DNAm variances in cancers compared to non-cancer samples [47, 48].

PCA of 7,484 samples from seven non-cancer tissues (adipose, nasal, blood, brain, buccal, sperm, and liver), which we also used to study DNAm variability (see below), showed clear clustering by tissue. The first two components explained 57% of total variance, with PC1’s contribution 38% and PC2’s contribution 19%. Sperm samples clustered far apart from the six somatic tissues (Figure 3c). After repeating PCA with sperm samples excluded, the first two components still explained over half (54%) of total variance, with PC1’s contribution 42% and PC2’s contribution 12% (Figure 3d).

### Over two-thirds of CpG probes that do not distinguish tissues map to gene promoters near CpG islands

CpG probes with low DNAm variation and low mean DNAm differences across experimental groups are less informative for quantifying group-specific DNAm differences. We analyzed autosomal DNAm variation in seven distinct tissues (adipose, nasal, blood, brain, buccal, sperm, and liver), as measured by noob-normalized, batch-corrected Beta-values (see Methods). We identified 4,577 probes each with consistently low variance (≤10th quantile) in each tissue and low difference between highest and lowest mean Beta-value (<0.01) across tissues (see Figures S7a and S8 as well as Table S7). Among probes with consistently low variance, 4,111 (90% of total) mapped to genes in CpG islands, typically at promoter regions of CpG island-overlapping genes (2,203 probes), and these fractions represented significant increases compared to the background of all autosomal CpG probes (binomial tests, p-values < 1e-3). It is likely the 4,577 probes are of low utility for quantifying DNAm differences across tissues, and their removal prior to performing an EWAS across non-cancer tissues could help increase statistical power.

### Over two-thirds of CpG probes that distinguish tissues map to genes

We identified 2,000 CpG probes in each of seven distinct non-cancer tissues (adipose, nasal, blood, brain, buccal, sperm, and liver) with high and tissue-specific variation in autosomal DNAm, as measured by noob-normalized, batch-corrected Beta-values (see Methods, Table S8, and Figures 4 and S7a). Distinctive patterns in DNAm across these probe sets may point to tissue-specific dynamics including cellular signaling or cell division rates. Compared to the background of all autosomal CpG probes, adipose and sperm had significantly lower fractions of gene-mapping probes, and all tissues except for blood had significantly greater fractions of both open sea-mapping probes and gene body-mapping probes (binomial tests, BH-adjusted p < 1e-3). Of the 14,000 total high-variance probes, 10,016 (71%) mapped to a gene region, typically at the gene body (8,006 probes, Table S8). The highest mean Beta-values were observed for nasal and adipose tissues (Figure 4a), and the highest variances were observed for sperm and adipose tissues (Figure 4b), while probes in blood had relative low means and variances. While most probes mapped to open seas in liver (1,014 probes), nasal (1,100), adipose (1,280), and sperm (1,063), greater fractions of open sea probes mapped to genes in liver (70% of open sea probes), nasal (74%), and adipose (71%) than in sperm (52%, Figure 4c).

**Figure 4.**
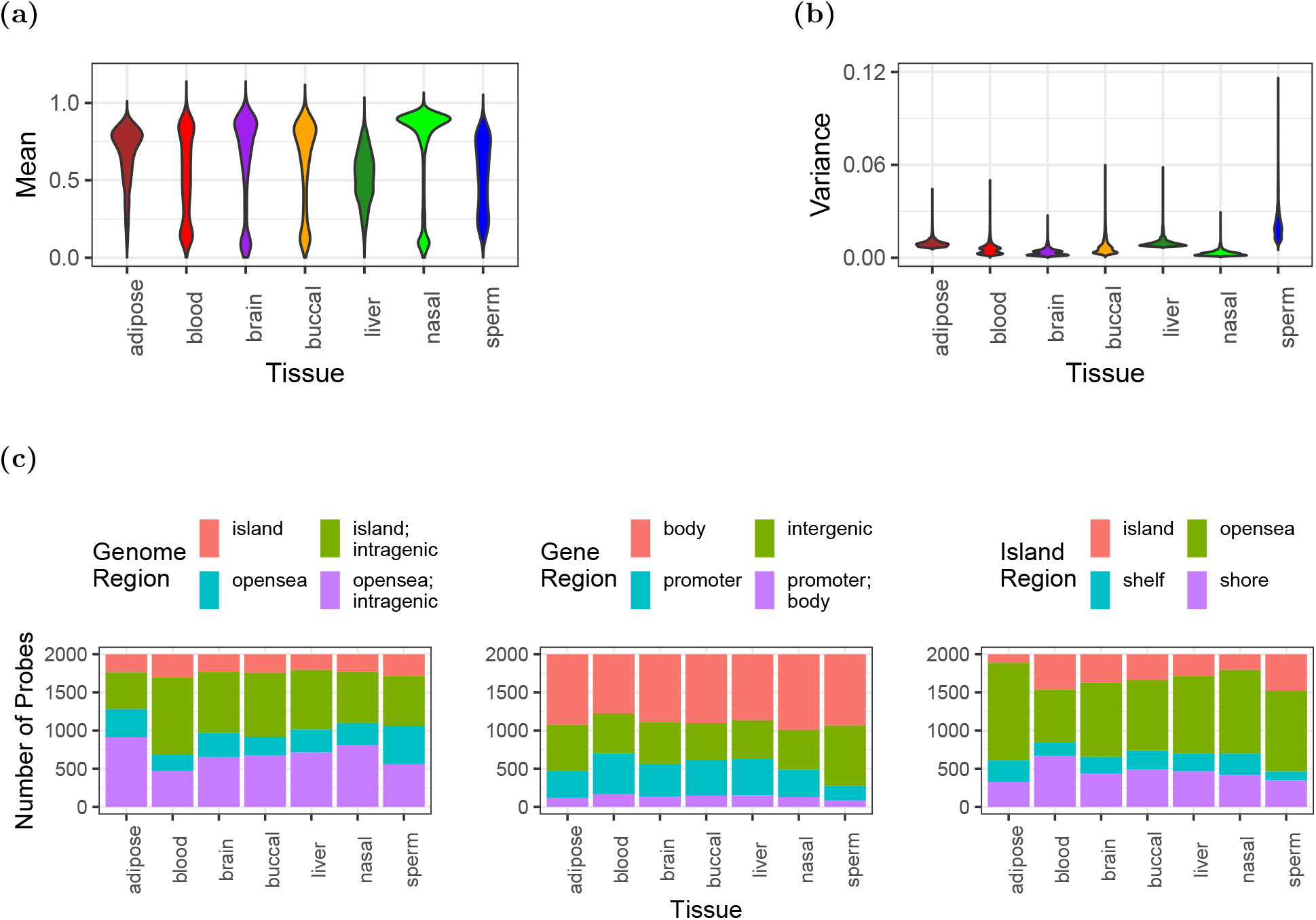
DNAm and genome mapping patterns among 14,000 CpG probes showing tissue-specific high variance in 7 tissues (2,000 probes per tissue, tissues: adipose, blood, brain, buccal, liver, nasal, and sperm). (a and b) Violin plots of Beta-value (a) means and (b) variances at tissue-specific probes. (c) Stacked barplots of genome region mappings (Number of CpG probes, y-axis) across tissue-specific probes (x-axis). Color fills depict (left) island and gene overlap, (center) gene region overlap, and (right) island region overlap.

### Normalized Beta-values for GEO DNAm array studies are rapidly accessed via the recountmethylation package

To accommodate a wide range of analysis strategies, DNAm assays and sample metadata were compiled into databases in two distinct formats, including Hierarchical Data Format 5 (HDF5) and `HDF5-SummarizedExperiment. HDF5-SummarizedExperiment` compilations are tailored for rapidly executing data summaries and query operations in the R/Bioconductor framework via `DelayedArray` objects. Raw red and green signals are provided as HDF5 (120 GB) and `HDF5-SummarizedExperiment` (119 GB) files, and raw methylated and unmethylated signals and noob-normalized Beta-values are provided as `HDF5-SummarizedExperiment` files (94 GB and 133 GB, respectively). The `recountmethylation` R/Bioconductor package facilitates database access as described in the User’s Manual. It allows full database utilization with rapid queries on the provided sample metadata, including model-based estimates for sex, epigenetic age, and blood cell types [5, 32, 33]. The package Data Analyses vignette further provides code to reproduce our comparisons of mined and epigenetic ages, sample storage type quality comparisons, and tissue-specific DNAm variability analyses described above.

## Materials and Methods

### Discovery and download of DNAm array IDATs on GEO

We used the `esearch` function of Entrez Programming Utilities v10.9 to search for every HM450K sample published to GEO as of March 31, 2019 for which two `gzip`-compressed IDAT download URLs were available. We ultimately downloaded IDATs for 35,360 samples (GSMs). Search and download were performed using the script https://github.com/metamaden/recount-methylation-server/blob/master/src/server.py.

### Preprocessing of DNAm array IDATs on GEO

We preprocessed DNAm array IDAT pairs for 35,360 HM450K samples on GEO using the R/Bioconductor package minfi v1.29.3 [32], applying the normalized exponential out-of-band probe method (i.e., noob normalization) in the analysis pipeline at https://github.com/metamaden/rmpipeline. The noob normalization technique mitigates run-specific technical biases and precedes batch-and/or study-level normalization steps [31].

### Quality control results

We computed 19 quality metrics from red and green color channel signals for HM450K samples (Table S2). To obtain the 17 BeadArray controls, we referred to Illumina’s official documentation [37,38] as well as methods in the `ewastools` v1.7 package [29]. For our calculations, we used signal from the extension green control as background, and we used a denominator offset of 1 where it would otherwise be 0 (see Supplemental Information) [29]). These calculations were done with the script https://github.com/metamaden/recountmethylationManuscriptSupplement/blob/main/R/beadarray_cgctrlmetrics.R. We thereby obtained a binary matrix of outcomes across the 17 BeadArray controls, where pass = 1, and fail = 0, on which we performed PCA using the “prcomp” R function from the stats v4.0.2 R package. We then used ANOVAs to determine the variances explained by each control for each component, and we obtained stacked barplots of component variances with `ggplot2` (Figure S4). The script https://github.com/metamaden/recountmethylationManuscriptSupplement/blob/main/inst/scripts/figures/figS4.R reproduces our steps.

We subsequently computed array-wide *log_2_* median methylated and *log_2_* median unmethylated signals, as reproduced in the `recountmethylation` data analysis vignette at https://www.bioconductor.org/packages/release/bioc/vignettes/recountmethylation/inst/doc/recountmethylation_data_analyses.pdf.

### Obtaining sample metadata

Sample metadata was downloaded from GEO as GSE-level Simple Omnibus Format in Text (SOFT) files using the script https://github.com/metamaden/recount-methylation-server/blob/master/src/dl.py. From SOFT files, GSM-level metadata were extracted as JSON-formatted files. Study-specific metadata fields were filtered prior to learning sample annotations (see below). These steps were performed using the scripts at https://github.com/metamaden/recountmethylationManuscriptSupplement/tree/main/inst/scripts/metadata.

### Learning sample annotations

We took a partially automated approach to learn sample annotations from mined metadata (Table S1). Our annotations were inspired by those in `Marmal-aid` [28] and included disease/experiment group, age, and sex (Table S1, [28]). To learn labels, we first coerced SOFT-derived metadata into annotation terms, then used manually constructed regular expressions to extract new labels (see Supplemental Information).

### Learning sample type predictions

We learned additional metadata using the MetaSRA-pipeline (https://github.com/deweylab/MetaSRA-pipeline [49], Table S1, [50]). This pipeline uses natural language processing to map sample metadata to curated ontology terms from the ENCODE project. It returns mapped terms and sample type confidences for each of six categories. We retained categories with the highest-confidence predictions as the most-likely sample types (see Table S1, Figure S2, and Supplemental Information).

### Model-based metadata predictions from DNAm

After noob normalization, we performed model-based predictions of sample age [5], sex [32], and blood cell type fractions [33] using the `minfi` (v.1.29.3) and `wateRmelon` (v.1.28.0) R/Bioconductor packages in our script https://github.com/metamaden/recountmethylationManuscriptSupplement/blob/main/inst/scripts/metadata/metadata_model_predictions.R. We tested concordance of mined and predicted sex and age to inform the use of these predictions and reliability of learned annotations (see Results).

### Principal component analyses of autosomal DNAm

We performed array-wide approximate principal component analyses (PCA) with the `stats` (v.3.6.0) R package, using noob-normalized autosomal DNAm from all samples and a subset of filtered samples from seven non-cancer tissues (Beta-values, Figures 3 and S6). Missing values were imputed by array-wide DNAm medians (Beta-value scale) within samples. To improve computational efficiency, we first applied feature hashing (also known as the hashing trick) [51,52] to project the normalized Beta-value arrays into an intermediate reduced space before performing PCA. PCA results were visually almost identical whether we invoked an intermediate dimension of 1,000 or 10,000 (results not shown). We used the 1,000-dimension mapping for analyses in Figure 3 (data provided in Supplemental Files). The above analysis steps are shown in the script https://github.com/metamaden/recountmethylationManuscriptSupplement/blob/main/inst/scripts/analyses/pca_analysis_fig3.R.

### Annotation of studies for cross-tissue DNAm variability analyses

We identified samples of seven distinct tissues (adipose, blood, brain, buccal, liver, nasal, and sperm), where each tissue included at least 100 samples across at least 2 GSE records (Table S6). We summarized study characteristics, including phenotype or disease of interest, in Table S6. Targeted samples were from a variety of studies targeting various diseases, syndromes, disorders, and exposures. Patient demographics spanned all life stages, including fetal, infant, child, and young and old adult, and several studies focused on ethnic groups not commonly studied (e.g. Gambian children from GSE100563;GSE100561).

### Preprocessing and analyzing seven non-cancer tissues for DNAm variability analyses

We studied samples in seven tissue types, including adipose, blood, brain, buccal, nasal, liver, and sperm (Table S6, Figures S7a and S7b). We removed likely low-quality samples that showed low study-specific (< 5th quantile) methylated and unmethylated signal, or showed signal below manufacturer-prescribed quality thresholds for at least one BeadArray control. We also removed putative replicates according to genotype-based identity predictions from `ewastools` (Supplemental Information, [29]).

We preprocessed noob-normalized DNAm for each tissue separately. First, we performed linear model adjustment on study IDs using DNAm M-values, defined as *logit(β*), with the `limma` v3.39.12 package. We then converted the adjusted DNAm to Beta-value scale. To account for the impact of confounding variables, we removed probes whose DNAm variances showed significant (p-adjusted < 0.01) and substantial (percent variance ≥ 10%) contributions from model-based predictions of age, sex, and cell type fractions, which removed 39,000 to 194,000 (8% to 40% of) probes across tissues (ANOVAs, Figure S7c).

After preprocessing, we identified probes with recurrent low variance and low mean intervals (max – min, mean tissue-wise DNAm, <0.01 or 1%) across seven distinct tissues. We also identified probes with high and tissue-specific variance. For each analysis we used a two-step probe selection process in each tissue where we selected (i) probes in the highest or lowest 10th quantile of variance (e.g. an absolute quantile variance filter), and (ii) probes in the highest or lowest 10th quantile variance across mean DNAm bins (e.g. a binned quantile variance filter, 10 bins of magnitude 0.1 or 10% DNAm, Figure S7a). The `recountmethylation` Data Analyses vignette reproduces these analyses for two tissues, and the full analysis scripts are contained at https://github.com/metamaden/recountmethylationManuscriptSupplement/tree/main/inst/scripts/analyses.

### Statistical analyses and visualizations

Statistical analyses and visualizations were conducted with the R and Python programming languages. We used the `numpy` (v1.15.1), `scipy` (v1.1.0), and `pandas` v0.23.0 Python packages to manage jobs and downloads, perform data extraction, and calculate summary statistics. We used the `minfi` v1.29.3 and `limma` v3.39.12 R/Bioconductor packages for downstream quality control, preprocessing, and analyses. Plots were generated using base R functions, `ggplot2` (v3.1.0), and `ComplexHeatmap` (v1.99.5) [53, 54]. To reproduce analyses, see Supplemental Methods and files at https://github.com/metamaden/recountmethylationManuscriptSupplement, and see the Data Analyses vignette in the `recountmethylation` R/Bioconductor package.

### Supplemental Information

Supplemental Information, including methods, code, scripts, and data files are accessible at https://github.com/metamaden/recountmethylationManuscriptSupplement. Large supplemental data files are accessible at https://recount.bio/data/recountmethylation_manuscript_supplement/.

### Companion R/Bioconductor package

Databases of the samples compiled and analyzed in this manuscript are accessible, along with comprehensive instructions and analysis examples, in the recountmethylation R/Bioconductor package at http://bioconductor.org/packages/devel/bioc/html/recountmethylation.html.

## Discussion

### Limitations of this study

We conducted a cross-study analysis of methylation array samples comprising a large subset of available HM450K samples on GEO. While we omitted studies using the HM27K and EPIC platforms, our compilation strategy could also be generalized to these platforms. Further, while our results suggest BeadArray controls could be improved by applying different quantitative thresholds for failure, it remains unclear whether a single universal threshold or multiple experiment-specific thresholds is desirable for each (Results, Figure S3). Nonetheless, five of 17 BeadArray controls (both Biotin Staining controls, both Non-polymorphic controls, and Bisulfite Conversion I Red) are demonstrably useful for assessing the quality of an experiment. We lacked a definitive gold standard set of well-described DNAm array samples, which would have allowed for more detailed estimations of metadata errors beyond direct concordances. Finally, our DNAm variability study across seven distinct non-cancer tissues used a within-tissue preprocessing approach. We were constrained this way because study-specific variation was high relative to tissue-specific variation, and effective study-and-tissue normalization would have required considerably more data from studies of multiple tissues than were available at the time of analysis.

### Recommendations for metadata reporting

Thoroughly characterizing samples submitted to public archives with accurate metadata makes them easier to repurpose for new studies. After manually inspecting hundreds of DNAm array studies on GEO, we formulated some best practices for the submitter who is labeling samples in a study to facilitate their discoverability and improve their utility for other investigators:

1. Include key attributes (sex, age, tissue, disease, etc.) even when any one is the same across a sample set, since that attribute may vary in a cross-study analysis.
2. Repeat GSE-level (i.e., study-level) metadata in GSM-level (i.e., sample-level) metadata. This includes sample types (e.g., tissue or cell line) and characteristics (e.g., storage conditions or preparation steps).
3. Include units of numerical variables to ensure their proper interpretation.
4. Clarify circumstances under which attributes were obtained where appropriate. Examples: age *at diagnosis, tumor-adjacent* normal tissue, blood *from leukemia patient.*

### Concluding remarks

We performed extensive analyses of 35,360 HM450K samples with IDATs from 362 studies in GEO, approximately three times the number of samples considered in prior cross-study analyses [5,28,29]. We further released the R/Bioconductor package `recountmethylation` including our new tissue and disease state labels, model-based predictions for age, sex, and blood cell composition, as well as noob-normalized Beta-values for array samples. This resource should prove valuable for reusing publicly available methylation data.

## Acknowledgments

This research was funded in part by NIH 5R01GM121459-02 to KH. We are especially grateful to John Greally for supportive comments and helpful suggestions as this work progressed. We also thank Julianne David, Mary Wood, Ben Weeder, Austin Nguyen, and Chris Loo for early feedback on our manuscript.

## Supporting Information

**Figure S1.**
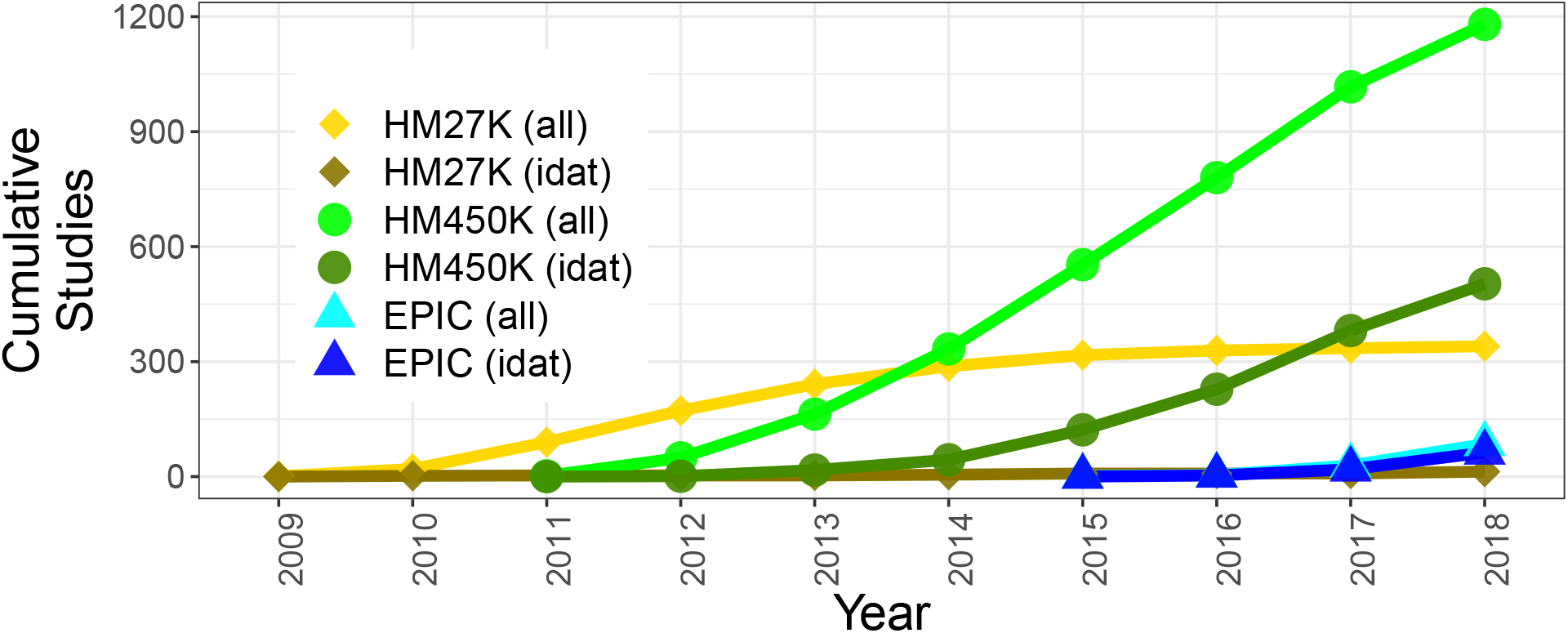
Cumulative yearly DNAm array studies in GEO for three platforms: HM27K (diamonds, yellow is all studies, brown is studies with IDATs), HM450K (circles, light green is all studies, dark green is studies with IDATs), and EPIC/HM850K (triangles, light blue is all studies, and dark blue is studies with IDATs).

**Figure S2.**
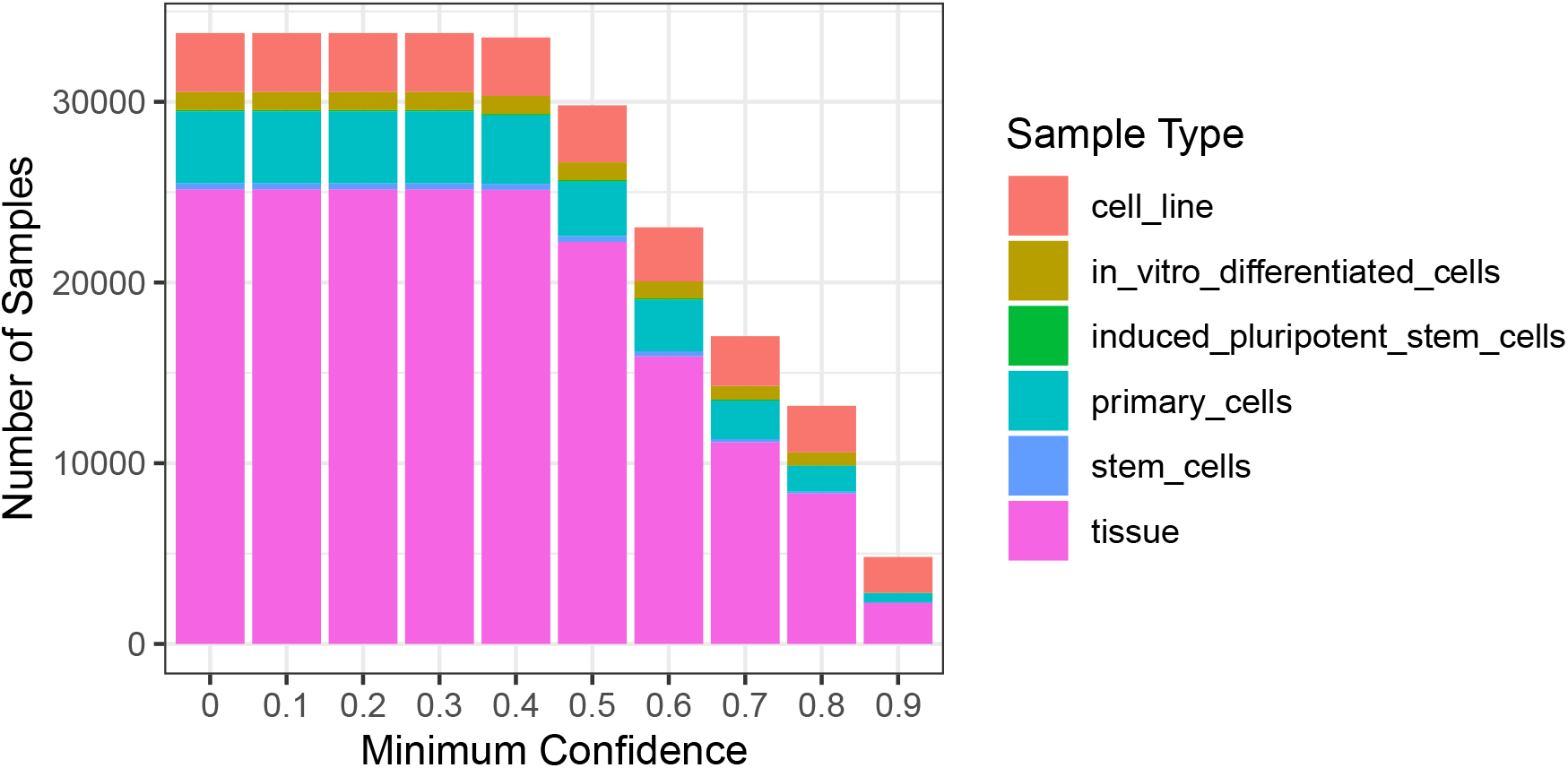
Most likely sample types (colors) predicted from MetaSRA-pipeline, by maximum confidence cutoffs (x-axis, barplots).

**Figure S3.**
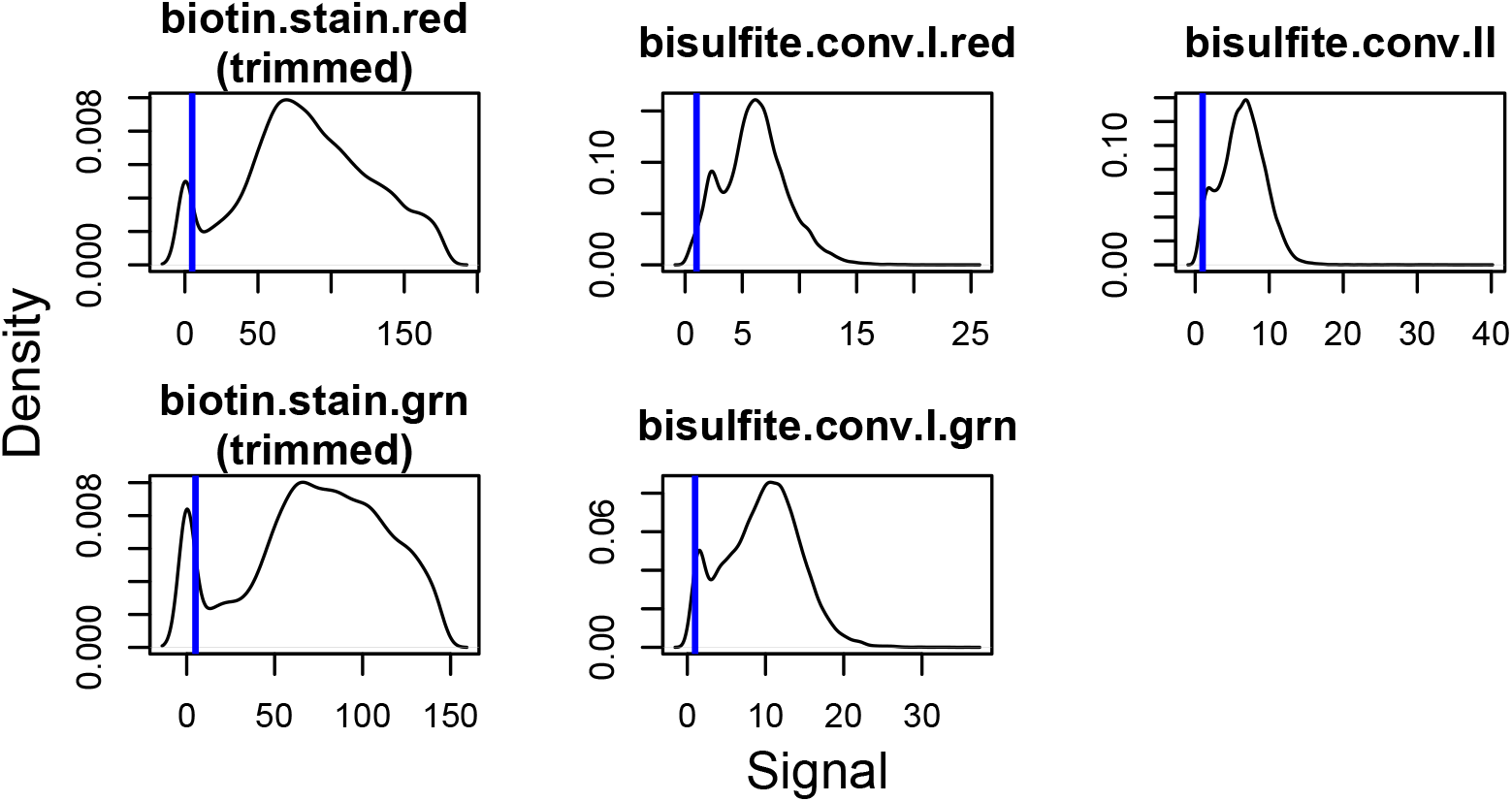
Signal density plots across five Illumina BeadArray controls, showing trimmed sample frequencies (x-axis is signal, y-axis is density) and published minimum signal thresholds (vertical blue lines).

**Figure S4.**
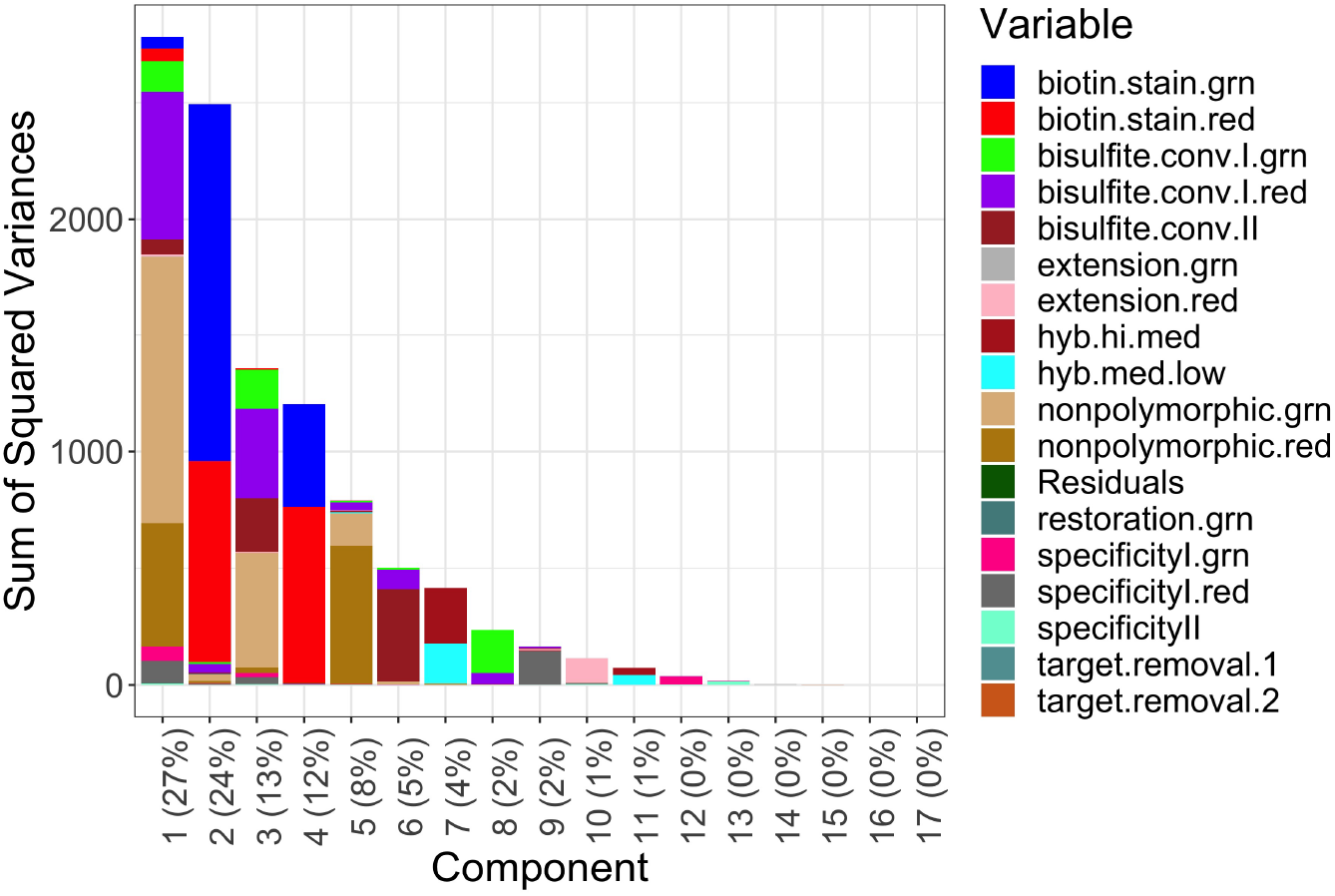
Screeplot from PCA of BeadArray control pass status (i.e., pass = 1, fail = 0) across samples, where the y-axis denotes the sum of the squared variance, the x-axis denotes principal component. Percentages in labels are corresponding component contributions to total variance. Colors capture magnitudes of component variances explained by each model variable from analyses of variance.

**Figure S5.**
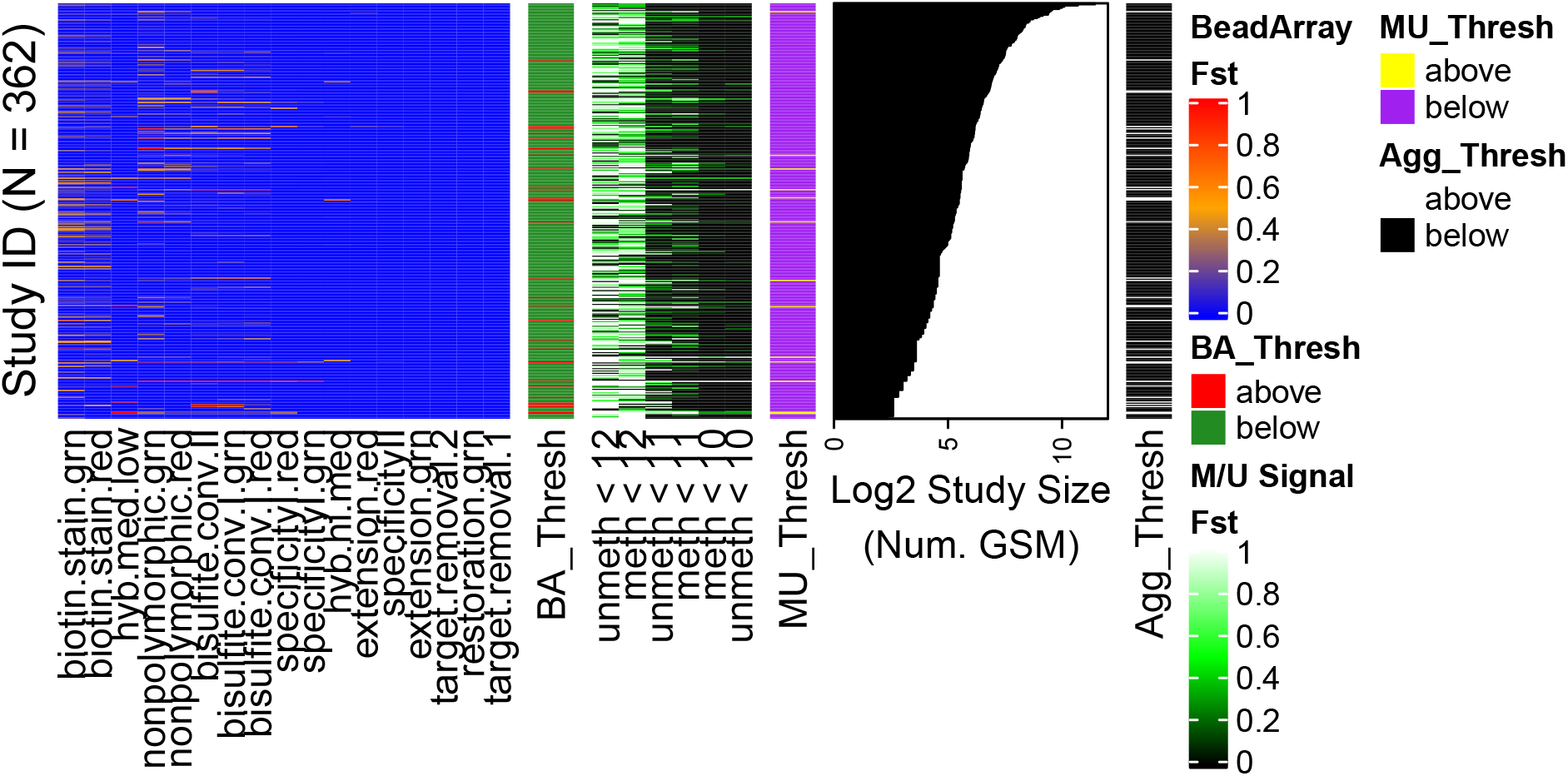
Heatmap of sample sub-threshold frequencies (*f_st_*) by study across quality metrics (BeadArray metric published thresholds, left, methylated [M] and unmethylated [U], *log_2_* median scale, middle). Annotations (small bars and far right barplots) are for whether a study showed >0.6 frequency of samples below at least one BeadArray metric(far left annotation, red, green is ≤ 0.6), whether a study showed >0.6 samples with ≤ 11 M and U signal (yellow, purple is <0.6, middle annotation), whether a study was above the sub threshold frequency for either BeadArray metric or signal annotation (right annotation, white, black is neither), and the *log_2_* scaled study size (far right barplots, number of GSMs), respectively. Only studies with data from >4 GSM IDs were included.

**Figure S6.**
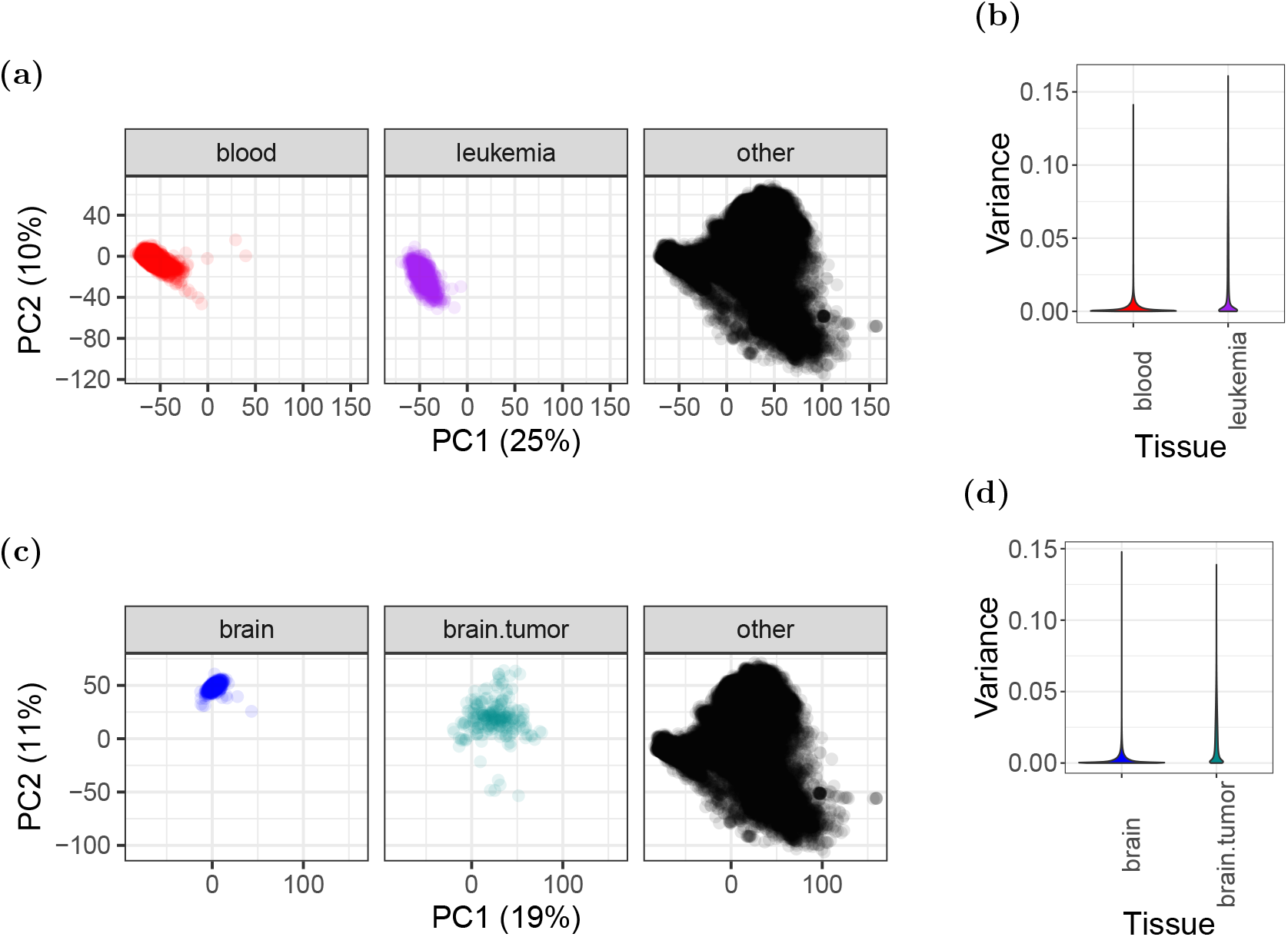
PCA facet plots and probe variance distributions of autosomal DNAm (noob-normalized Beta-values). (a) Facet plots of non-cancer blood (left, red), leukemias (middle, purple), and remaining samples (right, black) from Figure 3a. (b) Violin plots of DNAm variances in blood and leukemia samples. (c) Facet plots of non-cancer brain (left, blue), brain tumors (middle, dark cyan), and remaining samples (right, black) from Figure 3b. (d) Violin plots of DNAm variances in brain and brain tumor samples.

**Figure S7.**
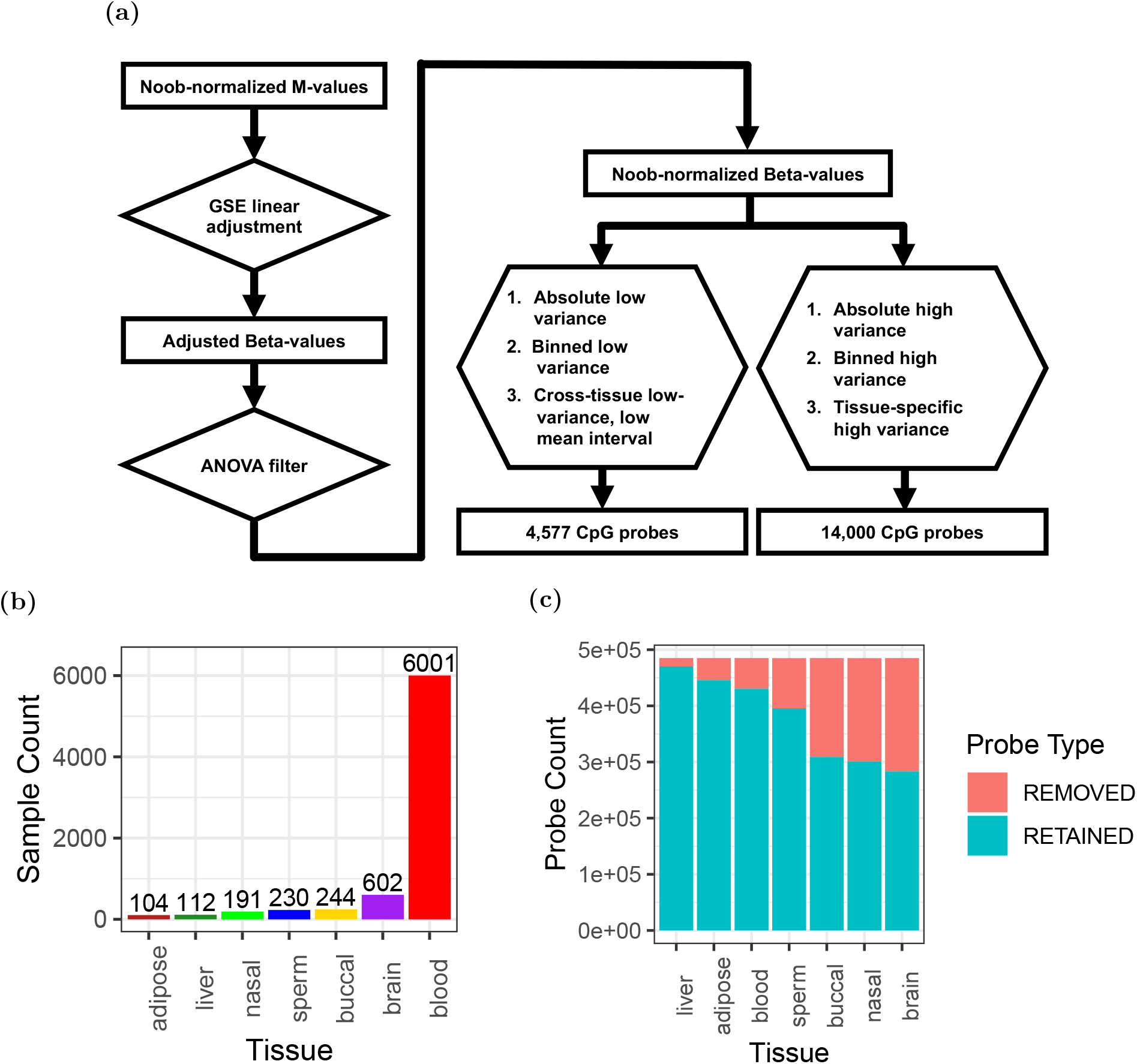
Analysis of autosomal DNAm variation across 7 non-cancer tissues (adipose, blood, brain, buccal, nasal, liver, and sperm). (a) Workflow to normalize, preprocess, and analyze DNAm variation within 7 tissues (adipose, buccal, brain, liver, sperm, nasal, and blood), including references to relevant figures. (b) Numbers of available samples by tissue type, with number labels showing barplot amounts. (c) Number of probes removed and retained from ANOVA filters within tissues (see Methods).

**Figure S8.**
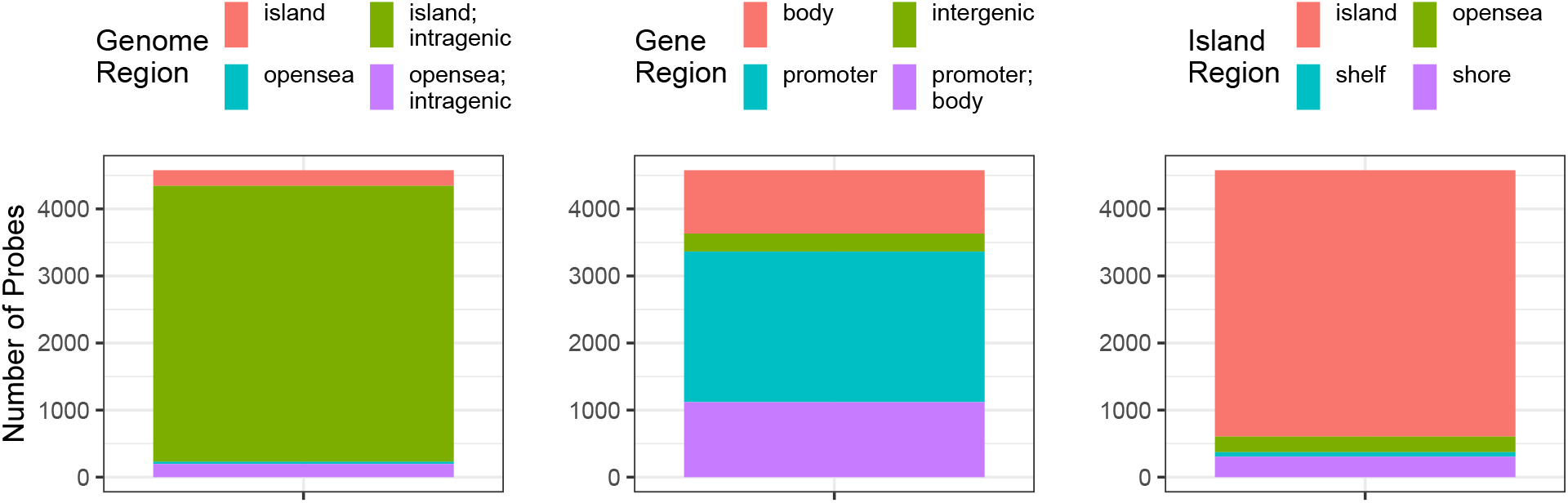
Genome mappings among 4,577 CpG probes with low variance across 7 tissues (adipose, brain, buccal, nasal, blood, liver, and sperm, see Methods, Figure S7). Color fills depict (left) Island and gene region overlaps, (middle) gene region overlaps, and (right) island region overlaps.

