## Supplemental Methods for "Human methylome variation across Infinium 450K data on the Gene Expression Omnibus"

### Supplemental Information for *Human methylome variation across Infinium 450K data on the Gene Expression Omnibus*

**Works cited** **4**

#### Overview

This document contains supplemental information for the manuscript *Human methylome variation across Infinium 450K data on the Gene Expression Omnibus*, including details about data access, aggregation, and analyses, with links to specific scripts where applicable.

#### Gene Expression Omnibus (GEO) data

DNA methylation (DNAm) array samples were identified as published to the Gene Expression Omnibus (GEO) and available in the GEO Data Sets database as of March 31, 2019.

##### GEO queries and data summaries

The Entrez Utilities software v10.9 was used to quantify DNAm array sample availability by year and platform for 3 major Illumina BeadArray platforms (HM450K, HM27K, and EPIC/HM850K).

Samples and studies were identified for download using the script https://github.com/metamaden/recount-methylation-server/blob/master/src/edirect_query.py.

Data availability by year was determined using the script https://github.com/metamaden/recountmethylationManuscriptSupplement/blob/main/inst/python/eqplot.py.

Plots for figures 1A and S1 were generated using the scripts https://github.com/metamaden/recountmethylationManuscriptSupplement/blob/main/inst/scripts/figures/fig1a.R and https://github.com/metamaden/recountmethylationManuscriptSupplement/blob/main/inst/scripts/figures/figS1.R.

#### Data acquisition

We used the Python programming language to develop a download management system to handle and version file downloads from GEO. We used Celery v5.0.0 to handle job management and logging. We used this job management system to obtain GSM IDATs, GSE SOFT files, and other data using batch queries to the GEO Data Sets repository. IDATs and SOFT files were acquired using the script https://github.com/metamaden/recount-methylation-server/blob/master/src/dl.py and download jobs were managed using the script https://github.com/metamaden/recount-methylation-server/blob/master/src/server.py

#### Processing DNAm data from IDATs

Signals were read from sample IDATs into an R session using the `minfi` v1.29.3 R/Bioconductor package (Aryee et al. 2014). DNAm assay data were read from IDATs using the script https://github.com/metamaden/rmpipeline/blob/master/R/rmpipeline.R.

### Sample metadata

Scripts for metadata processing can be found in the directory https://github.com/metamaden/recountmethylationManuscriptSupplement/tree/main/inst/scripts/metadata

#### Extracting sample/GSM metadata from GSE SOFT files in JSON format

After obtaining the GSE SOFT files from GEO, we extracted GSM-specific metadata as JSON-formatted files using the script https://github.com/metamaden/recountmethylationManuscriptSupplement/tree/main/inst/scripts/metadata/soft2json.R. Before further processing, we filtered the sample JSON-formatted metadata using the script https://github.com/metamaden/recountmethylationManuscriptSupplement/tree/main/inst/scripts/metadata/jsonfilt.R. This step attempted to remove any GSE-specific metadata while retaining sample-identifying characteristics.

#### Processing the sample metadata

The filtered JSON metadata were read into a list of tables, organized by GSE ID, using the script https://github.com/metamaden/recountmethylationManuscriptSupplement/tree/main/inst/scripts/metadata/make_gse_annolist.R. We then preprocessed these tables by moving the available metadata under common variables as shown in the script https://github.com/metamaden/recountmethylationManuscriptSupplement/tree/main/inst/scripts/metadata/metadata_preprocessing.R. This step was coordinated manually and separately for each respective GSE ID. We additionally obtained GSM record titles from JSON files using the script https://github.com/metamaden/recountmethylationManuscriptSupplement/tree/main/inst/scripts/metadata/get_gsm_titles.R. We postprocessed available metadata by mapping available data to controlled term vocabularies using the script https://github.com/metamaden/recountmethylationManuscrip

tSupplement/tree/main/inst/scripts/metadata/metadata_preprocessing.R. Controlled vocabularies were heavily inspired by terms used by the Marmal-aid resource, and we attempted to map the most frequently available and informative metadata we observed in the original SOFT files.

##### Predicting sample types from filtered sample JSON files

We ran the MetaSRA-pipeline software on the filtered versions of sample JSON metadata files (Bernstein et al. 2017). For this, we used a forked the original software available at https://github.com/metamaden/MetaSRA-pipeline. For each sample record, the MetaSRA-pipeline mapped a series of curated ontology terms and predicted the likelihoods for each of 6 sample type categories (Figure S2). We extracted the pipeline-mapped files into tables using the script https://github.com/metamaden/recountmethylationManuscriptSupplement/tree/main/inst/scripts/metadata/get_msrap_mdmap.R. We retained the most likely predicted sample types and their likelihoods as shown in the script https://github.com/metamaden/recountmethylationManuscriptSupplement/tree/main/inst/scripts/metadata/make_md_final.R.

##### DNAm model-based predictions for age, sex, and blood cell types

Model-based predictions of sex, age, and cell type fractions were obtained from DNAm data as described in https://github.com/metamaden/recountmethylationManuscriptSupplement/tree/main/inst/scripts/metadata/metadata_model_predictions.R. Age predictions were obtained by running the `agep` function from the `wateRmelon` v1.28.0 package on noob-normalized DNAm Beta-values (Horvath 2013). Sex predictions were obtained by running the `getSex` function from `minfi` on genome-mapped `MethylSet` data (Aryee et al. 2014). Cell fraction predictions were obtained by running the `estimateCellCounts` on `RGChannelSet`-formatted data (Houseman et al. 2012).

#### Quality and summary metrics

We calculated quality signals for 17 BeadArray controls, methylated and unmethylated signals, and likely sample replicates by study, as described in the table script https://github.com/metamaden/recountmethylationManuscriptSupplement/tree/main/inst/scripts/tables/tableS2.R. BeadArray signals for 17 controls were calcuated using the script https://github.com/metamaden/recountmethylationManuscriptSupplement/tree/main/R/beadarray_cgctrlmetrics.R. Our method was developed in consultation with Illumina's platform documentation and functions from the `ewastools` R package ("Illumina Genome Studio Methylation Module V1.8" 2010; "BeadArray Controls Reporter Software Guide" 2015; Heiss and Just 2018). Sample genetic identities were predicted using the functions `call_genotypes()` and `check_snp_agreement()` from the `ewastools` R package, which uses high-frequency SNP signals from HM450K arrays (Heiss and Just 2018). We predicted and recorded likely genotype-based replicates within each GSE record.

#### Statistical analyses

##### Tests, summaries, and plots

Sample data were obtained, read, and analyzed programmatically using the R v4.0.0 and Python v3.7.0 languages in a CentOS 7 remote server environment. IDAT signals were read into `SummarizedExperiment` objects using the `minfi` package. Summary statistics were generated using base R functions. Scripts to reproduce manuscript analyses are stored at https://github.com/metamaden/recountmethylationManuscriptSupplement/tree/main/inst/scripts/analyses. Statistical tests were performed using the `stats` v4.0.0 R package. Correlation tests used the Spearman method by setting `method = "spearman"` in the `cor.test` function. Analyses of variance (ANOVAs) were performed using the `anova` function. Label enrichments were tested using Binomial with the `binom.test` function, and T-tests used the `t.test` function. Unless noted otherwise, p-value adjustments used the Benjamini-Hotchberg method by setting `method = "BH"` in the `p.adjust` function. Plots used base R functions and the R packages `ggplot2` v3.1.0 and `ComplexHeatmap` v1.99.5. Scripts to reproduce manuscript figures are stored at https://github.com/metamaden/recountmethylationManuscriptSupplement/tree/main/inst/scripts/figures.

##### PCAs of autosomal DNAm

Approximate array-wide PCA was performed using noob-normalized Beta-values from autosomal probes and the `prcomp` from the `stats` R package. We condensed autosomal DNAm across 35,360 samples into 1,000 hashed feature columns, the performed cluster analysis. As a dimensionality reduction step, feature hashing or the "hashing trick" mapped CpG probe data into a smaller intermediate dimension while approximately preserving between-sample variation (Weinberger et al. 2010). Feature hashing was performed as shown in the script https://github.com/metamaden/recountmethylationManuscriptSupplement/tree/main/inst/python/dnamhash.py. The hashed features data is available at https://recount.bio/data/recountmethylation_manuscript_supplement/data/pca_fh1k_all_gsm35k.zip.
